# A multiscale framework for disentangling the roles of evenness, density and aggregation on diversity gradients

**DOI:** 10.1101/851717

**Authors:** Daniel J. McGlinn, Thore Engel, Shane A. Blowes, Nicholas J. Gotelli, Tiffany M. Knight, Brian J. McGill, Nathan Sanders, Jonathan M. Chase

## Abstract

Disentangling the drivers of diversity gradients can be challenging. The Measurement of Biodiversity (MoB) framework decomposes changes in species diversity into three components of community structure: the species abundance distribution (SAD), the total community abundance, and the within-species spatial aggregation. Here we extend MoB from categorical treatment comparisons to quantify variation along continuous geographic or environmental gradients. Our approach requires sites along a gradient, each consisting of georeferenced plots of abundance-based species composition data. We demonstrate our method using a case study of ants sampled along an elevational gradient of 28 sites in mixed deciduous forest of the Great Smoky Mountains National Park, USA. MoB analysis revealed that ant species richness decreased along the elevational gradient because of changes in the SAD and in spatial aggregation, but not because of changes in the number of individuals. Specifically, with increasing elevation, species evenness was lower and species were less aggregated. These results do not support the more-individuals hypothesis; alternative hypotheses are required to explain why evenness and aggregation decrease with elevation. Our extension of MoB has the potential to elucidate the drivers of diversity along environmental gradients and should be useful for a variety of assemblage-level data collected along gradients.

## Introduction

A critical limitation of most studies examining patterns of biodiversity along ecological or biogeographic gradients is that the most common measure of biodiversity--species richness--is limited in its utility for differentiating between several competing hypotheses that contribute to spatial variation in biodiversity. This limitation arises for two related reasons: (1) species richness is a summary variable of the relative and absolute numbers of individuals in a population, as well as their spatial distribution; (2) species richness depends on spatial scale in a non-linear way (Rahbek, 2005; Chase *et al.*, 2018; McGlinn *et al.*, 2019).

Examining variation in the total and relative abundance, as well as the spatial distribution of species along environmental gradients provides information that allows for distinguishing among drivers of biodiversity. For example, species richness is typically a positive function of the amount of energy that enters an ecosystem. One prominent hypothesis for this relationship is that the energy input into an ecosystem leads to increases in the numbers of individuals, which in turn supports higher species richness (Wright 1983, Evans et al. 2008). Under this ‘more individuals hypothesis’ (Srivastava and Lawton 1998) changes in species richness would be expected to be closely linked to changes in total numbers of individuals but not changes in species relative abundances or their spatial distributions. In contrast, if higher energy decreased competitive exclusion then changes in richness could be linked to changes in the relative abundance of species rather than the total number of all individuals (Evans et al. 2005, Hurlbert and Jetz 2010). Additionally, if energy changes the spatial pattern or relevance of environmental heterogeneity then species spatial structure would be expected to change. As a result, data and analyses that explicitly incorporate abundances of species and their spatial distribution across scales, rather than just a single scale-agnostic measure, can provide deeper insights into the potential underlying causes of variation in biodiversity.

The Measurement of Biodiversity (MoB) framework (Chase *et al.*, 2018; McGlinn *et al.*, 2019) was developed to explicitly dissect the abundance and distribution patterns that underlie changes in species richness. Specifically, MoB decomposes spatial variation in richness into the contributions from three components of community structure:

1. species abundance distribution (SAD) (including evenness and the size of the species pool). A more even community will have more species found in a given sample than a less even community;
2. the community-level density of individuals (*N*); simply by sampling more individuals from a species pool, more species will be found;
3. intra-specific spatial aggregation. When species are aggregated (clumped) in the community, local species richness will typically be lower, while regional scale richness and beta-diversity can be higher.

These three components are largely sufficient for predicting many macroecological patterns of species richness (McGill 2010) and thus provide an important starting point for deciphering biodiversity patterns (see also He and Legendre 2002, Chase and Knight 2013). If species richness differs from one site to another, it does so because the SAD, N, and/or aggregation of species changes between those sites.

As it was originally developed (Chase et al. 2018, McGlinn et al. 2019), MoB consists of two complementary analyses for examining if a discrete explanatory variables (e.g., an experimental treatment like the presence or absence of a top predator) influences biodiversity: the two-scale, multimetric analysis and the multiscale, richness analysis. However, discrete variables are not the only variables that influence spatial variation in species richness. Species richness often varies along continuous gradients as well, such as gradients in temperature, latitude, or elevation. It is straightforward to extend the two-scale, multi-metric MoB to gradients using traditional regression analyses (e.g., generalized linear models) (e.g., Blowes et al. 2017, Chase et al. 2018). However, the multi-metric MoB only examines changes at discretely defined local (α) and regional (γ) scales. The multiscale MoB examines a range of scales, but it is less straightforward to extend to gradients.

Here, we outline an extension of multiscale MoB that allows for disentangling the drivers of species richness along continuous geographical or environmental gradients. We provide a conceptual overview and exposition within the mobr R package (McGlinn et al. 2019) to dissect the influence of the components of species richness (N, SAD, and aggregation) across ecological gradients. We apply the approach to a case study on spatial variation in ant diversity along an elevational gradient in the southern Appalachian mountains (USA)(from Sanders *et al.*, 2007). We demonstrate that the application of multiscale MoB helps to reveal how changes in the SAD, N, and intraspecific aggregation contribute to the multiscale pattern of richness change along gradients.

## Methods

To illustrate the motivation and the method of extending the multiscale MoB framework it is helpful to consider three simple scenarios (Fig. 1) in which only a single component of community structure is responsible for variation in species richness along a gradient. For example, richness may decline along a gradient due to a decrease in evenness (Fig. 1A, SAD effect), a decrease in the number of individuals (Fig. 1B, N effect), or increased intraspecific aggregation (Fig. 1C, aggregation effect). Although in reality, any change in species richness along any gradient is likely caused by changes in more than one of these components of community structure. Nevertheless, this simple example illustrates three key points: 1) species richness can change at one scale (plot scale) but not another (site scale), 2) species richness can change in a similar ways due to very different changes in the underlying components, and 3) a more direct focus on changes in these components and their contributions to changes in species richness across scales can elucidate the underlying causes of these changes.

**Figure 1.**
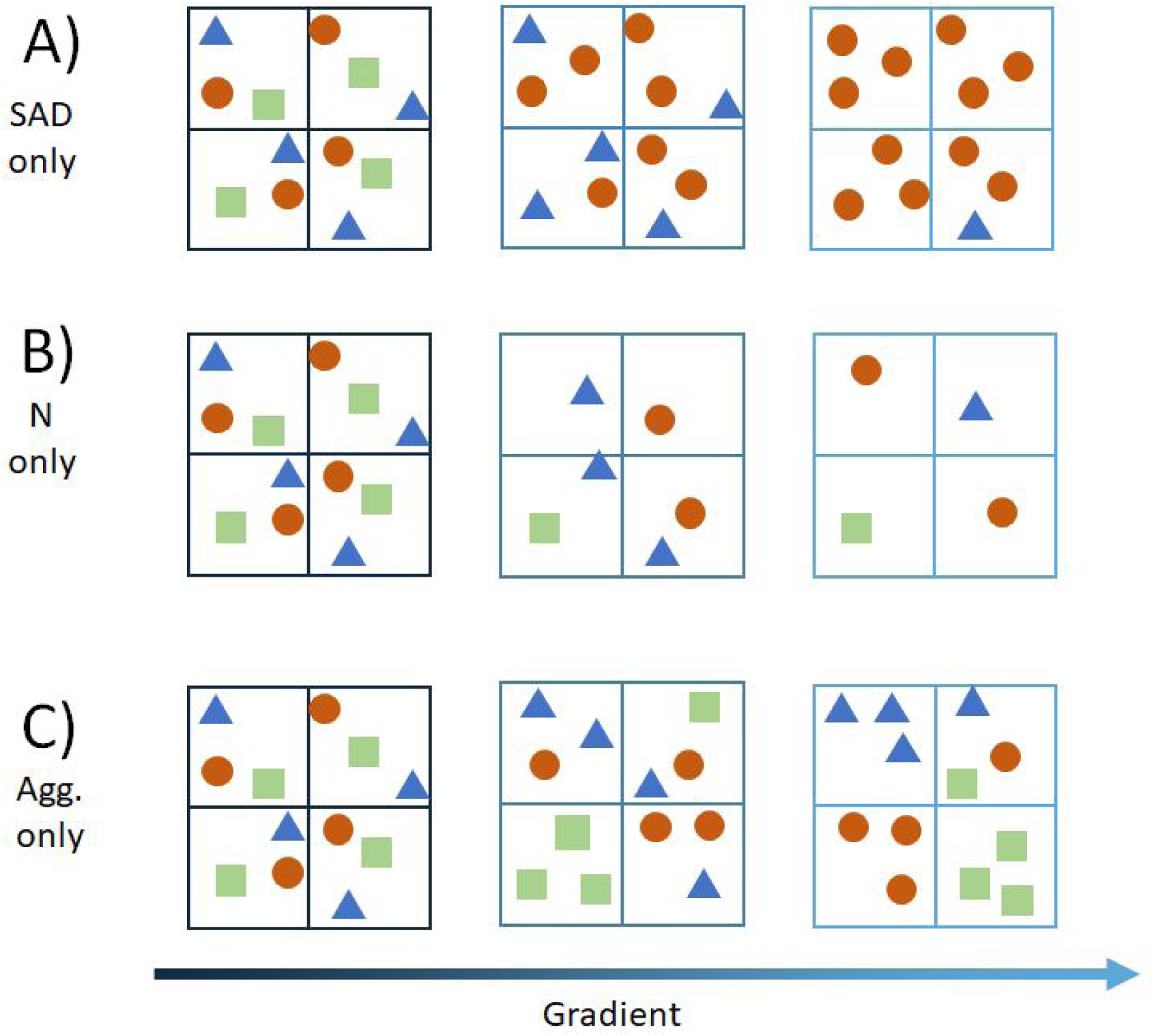
Cartoon communities from three sites arranged along a gradient (color gradient from dark blue to light blue) in three simple scenarios in which only the A) SAD, B) N, or C) aggregation shifts along the gradient. The large boxes represent sites, the small boxes represent plots, and the different symbols represent individuals of different species.

Although each of these scenarios results in a decrease in plot-scale species richness along the gradient, our extension of the multiscale MoB framework can disentangle which components of community structure are responsible for the change in *S* across scale (Fig. 2). Here we define scale as the number of samples (i.e., “plots”) or the number of individuals accumulated (McGill 2011). Multiscale MoB takes advantage of the unique information captured by three different types of rarefaction curves (Fig. 2):

**Figure 2.**
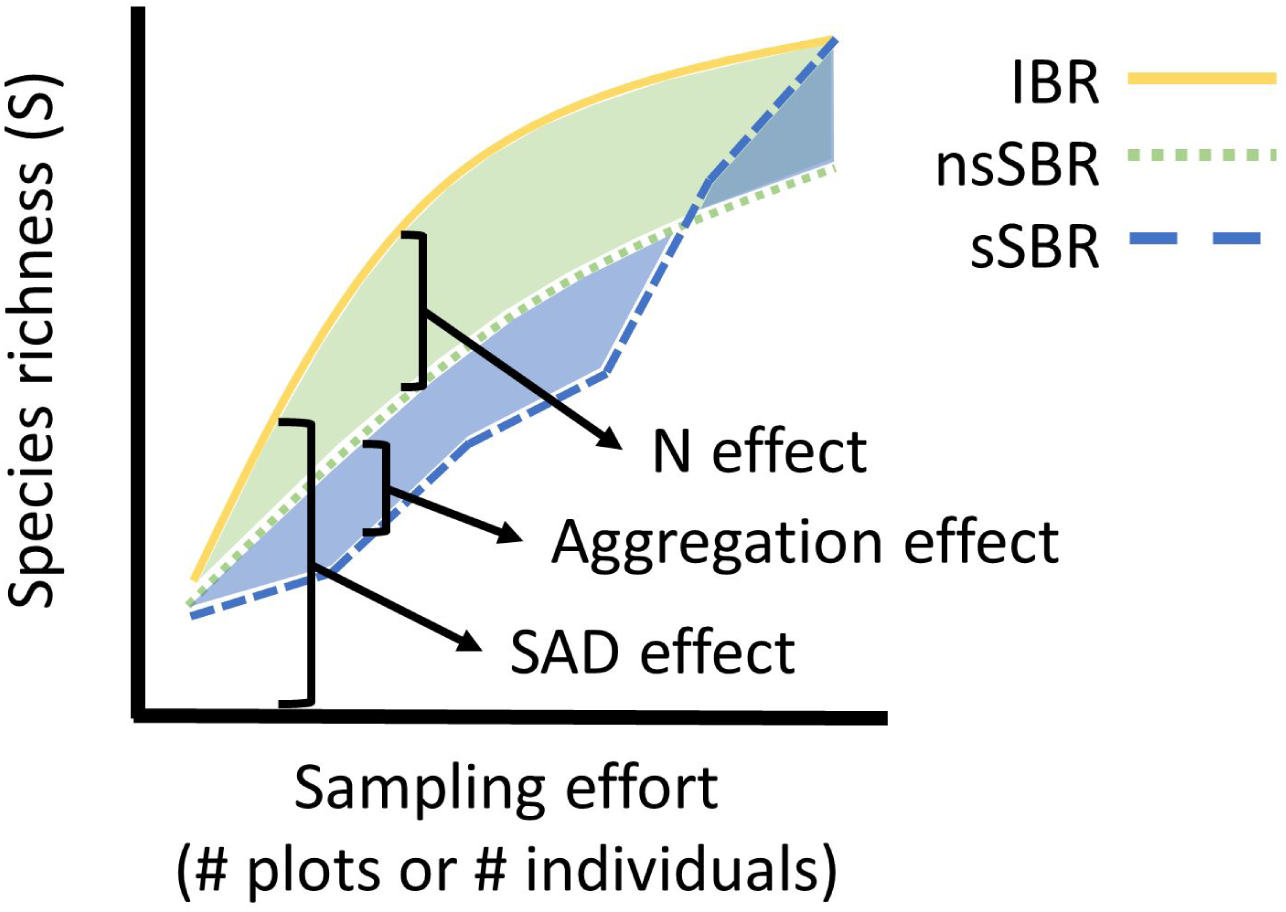
The three rarefaction curves compared at one site along a gradient in which this particular site has lower individual density than an average site on the gradient (i.e., a negative N effect is illustrated here). The individual-based rarefaction (IBR) is a direct expression of the SAD (yellow line). The non-spatial, sample-based rarefaction (nsSBR) reflects both the SAD and variation in N, thus the difference between the nsSBR and the IBR provides an estimate of the N effect (light green area). The spatial, sample-based rarefaction (sSBR) also takes spatial position into consideration, thus the effect of spatial aggregation is the difference between the sSBR and the nsSBR (light blue area). Note that the nsSBR must eventually intersect the IBR and sSBR at this site (i.e., all curves converge to the same total S once enough effort is considered).

- Spatial, sample-based rarefaction (sSBR) is the accumulation of species by collecting the closest plots first. All possible focal samples are considered and the resulting curves are averaged over (Fig. 2). The sSBR reflects information on aggregation, *N*, and the SAD.
- Non-spatial, sample-based rarefaction (nsSBR) is the number of species given *k* plots in which all *N* individuals are randomly re-assigned to plots while maintaining observed individual density (Fig. 2). The nsSBR reflects variation in both N and the SAD.
- Individual-based rarefaction (IBR) is the number of species given a random sample of *n* individuals out of N total individuals (Fig. 2). The IBR only reflects variation in the SAD.

Comparing each of these curves allows us to dissect the influence of the components on changes in *S* (Fig. 2). The difference between the sSBR and the nsSBR estimates the signature of aggregation on *S*; the difference between the nsSBR and the IBR reflects the influence of *N* on *S*; the IBR eliminates *N* and aggregation effects, and is just a reflection of the SAD (Fig. 2). In the context of analyzing changes in *S* along a gradient, the relevant question is how the effects of aggregation, *N*, and the SAD vary along the gradient across scales.

If only the SAD changes along the gradient (e.g., decreasing evenness in the illustrated scenario), the IBRs will diverge as sampling effort increases (Fig. 3A, gradient location represented by dark blue to light blue line colors, as in Fig. 1). Because the IBRs diverge, the strength of the detected SAD effect increases with effort (Fig. 3B). We can estimate the relationship between the gradient and the SAD effect on S using linear models (or non-linear if more appropriate) (Fig. 3B, only three scales shown for clarity) which allow us to quantify whether the strength of this relationship shows scale-dependence (Fig. 3C). If this component of community structure does not help to explain shifts in S along the gradient then its effect size (e.g., slope) will be zero (the dashed lines in Fig. 3C) and not different from a null model of random samples from the pooled across site SAD (described in McGlinn et al. 2019).

**Figure 3.**
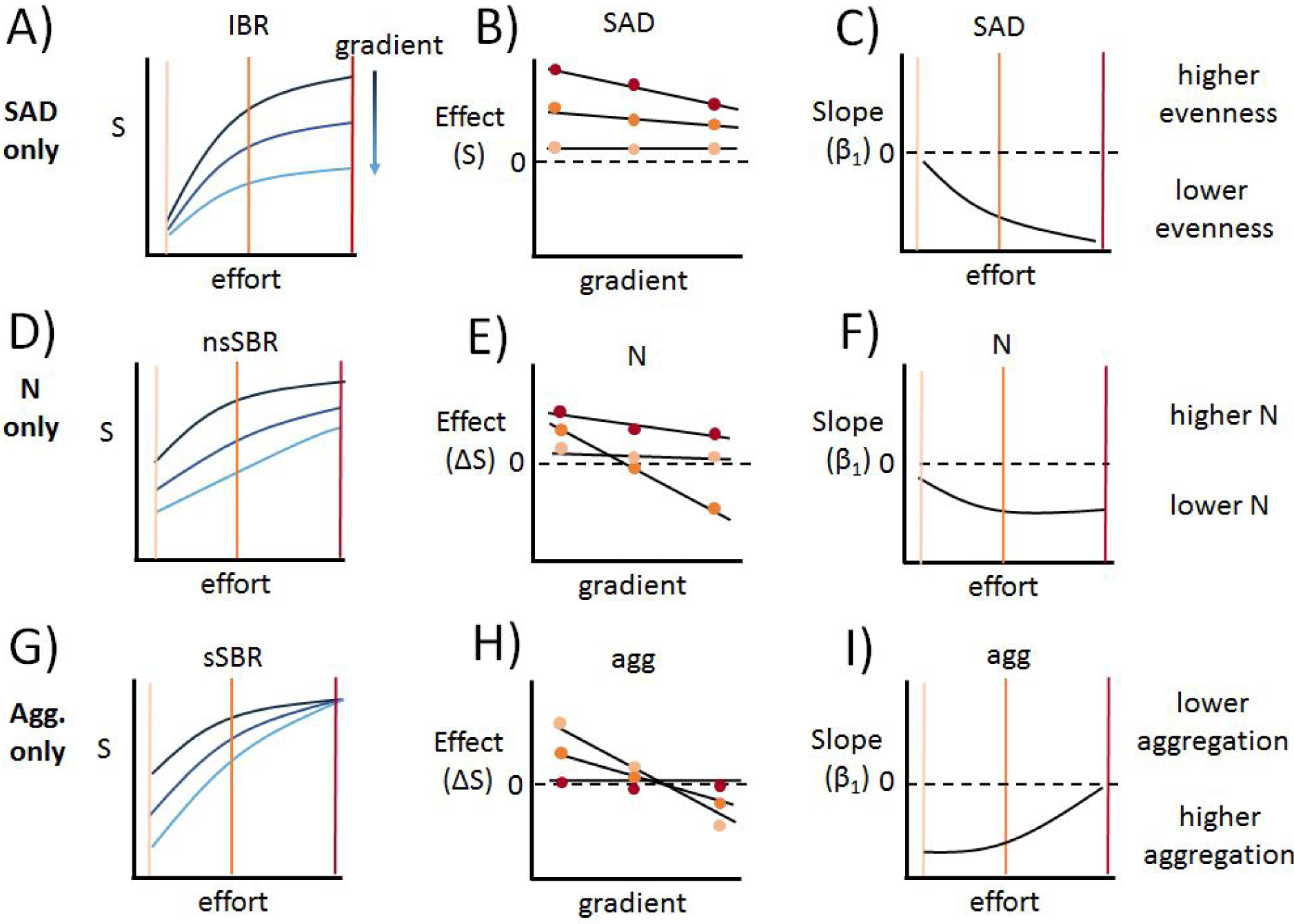
The three sets of hypothetical results illustrating the MoB multiscale approach using the cartoon communities considered in Fig. 1. Panels A, D, G display three types of rarefaction curves which detect different components of community structure (for clarity only the relevant rarefaction curves are shown to detect the component of community structure known to have shifted). IBR = individual-based rarefaction, nsSBR = non-spatial, sample-based rarefaction, and sSBR = spatial, sample-based rarefaction. For each type of rarefaction curve three curves are computed at each site along the gradient (colored dark blue to light blue as in Fig. 1). Three sampling efforts (orange vertical lines in panels A, C, D, F, G, I and points in B, E, H) are highlighted to emphasize that the variation in the curves (i.e., effect sizes) change with scale. Panels B, E, H display the strength of the SAD, N, and aggregation effects respectively on *S* plotted against the gradient. Regression lines are fit to the relationship between effect size and the gradient, and the strength (e.g., the estimated regression slope, β_1_) of those fits are plotted in panels C, F, I as a function of sampling effort. The dashed line denotes zero effect (B, E, H) or slope (C, F, I, the null expectation).

If only N changes across the gradient (e.g., decreasing N in the illustrated scenario), the nsSBRs vary along the gradient (Fig. 3D), but not the IBR and the sSBR curves (not shown). As with the SAD effects, we can model the relationship between the N effect (i.e., the difference between the nsSBR and IBR, Fig. 2) and the gradient across spatial grains (Fig. 3E). The net result is shown in Fig. 3F where the decrease in N along the gradient is captured as a negative slope.

Finally, if only species aggregation changes along the gradient, the sSBRs will vary along the gradient (Fig. 3G), but not the other two rarefaction curves (IBR and nsSBR). In our cartoon scenario, the strength of aggregation is strongest at fine spatial scales and decreases as the sSBRs converge (Fig. 3G, H, I). Note that regardless of the specific scenario considered in a balanced experimental design (i.e., same number of subplots at each site along the gradient), the effect of aggregation must converge on zero at the maximum sampling effort (i.e., all plots collected) because at this scale the sSBR must be identical to the nsSBR (McGlinn et al. 2019).

In summary, we have extended the multiscale MoB comparisons between categorical treatments to continuous gradients. This can be thought of as extending MoB from a t-test to a regression analysis. We have released a dev branch of the mobr R package on github (https://github.com/MoBiodiv/mobr/tree/dev) to carry out the following steps of the gradient analysis we described above:

1. Compute three rarefaction curves that carry with them different information on the influence of N, the SAD and aggregation for each set of samples (i.e., a site) along the gradient of interest: sSBR, nsSBR, and IBR (Fig. 3A, D, G).
2. Compute the differences between rarefaction curves of interest (sSBR - nsSBR and nsSBR - IBR) at each site; these are the estimates of effect size for aggregation and N respectively. (Fig. 3B, E, H).
3. Model the relationship between the gradient and the estimates of the SAD, N, and aggregation effects (Fig. 3B, E, H).
4. Examine how the model coefficients or fit vary with sampling effort. (Fig. 3C, F, I).
5. Compare the observed results to randomization-based null models (described in McGlinn et al. 2019) for each component of community structure (i.e., aggregation, N, and the SAD) (Fig. 3C, F, I) to infer if the effects and their relationship to the gradient are different than expected purely due to chance.

In our cartoon example, S decreases along the gradient, as it often does along environmental gradients. But, using the MoB approach, we can estimate which component of community structure - N, SAD, or aggregation - influence spatial variation in richness. Although the cartoon only demonstrated changes in richness due to a single component of community structure, it is likely that more than one component will be responsible for spatial variation in richness in real communities. A sensitivity analysis suggested that the multiscale MoB approach can reliably detect the signature of simultaneous changes in multiple components of community structure on richness as well (McGlinn et al. 2019).

### Data requirements

The cartoon in Fig. 1 illustrates the basic data requirements to use MoB to explore variation in S along gradients. Obviously, sampling sites must be distributed along an environmental gradient. At each sampling site, there must be a collection of several (≥5) geo-referenced samples that contain data on the abundances and identities of each species in a sample. It is not necessary for the sampling design to have the same number of samples at each site along the gradient, but the sSBR should be truncated to the smallest common number of samples per site across the gradient (to minimize any influence of spatial extent). Similarly, the IBR and the nsSBR should be truncated to the smallest number of individuals observed and therefore sites (not necessarily samples) should have enough individuals so that rarefaction results are meaningful - differences in rarefaction curves are constrained to be small at low sample sizes. It is also important that the spatial grain and spatial arrangement of plots is consistent along the gradient. Otherwise the investigator runs the risk that the variation in sampling design is responsible for changes in the components of community structure. If a given sampling design is not consistent among sites along a gradient, then it may be necessary to subset the samples so that sites along the gradient have comparable spatial extents. It is more important to ensure a constant extent across sites than a balanced design when using rarefaction curves to compare biodiversity. Although there may be slight differences in their numbers and spatial arrangement, it is more important that samples are standardised across all sites so they relate to a constant unit of effort (e.g. area).

### Case study

To demonstrate our new methods we use data from Sanders et al (2007) who examined spatial variation in richness along an elevational gradient (379–1742 m) in the Great Smoky Mountains National Park, USA. Sanders et al. (2007) collected ant samples from each site along the elevational gradient by visiting each site once between June–August in 2004-2006 when ants in the national park are typically most active (Dunn *et al.*, 2007b). All sites were located in mixed hardwood forests and away from any area of recent human disturbance. We removed one site (site code = “NODI”) which only contained 6 individuals across the 16 samples, resulting in a dataset of 28 sites.

At each site, data come from a randomly placed a 50 × 50 m plot, from which 16 1-m^2^ quadrats were arranged in a nested design: 10 × 10 m subplots were placed in the corners of each 50 × 50 m plot, and 1-m^2^ quadrats were placed in the corners of each 10 × 10 m subplot, for a total of 16 1-m^2^ quadrats per site. Ants were sampled by collecting all of the leaflitter within each quadrat and sifting through it with a coarse mesh screen (1-cm grid) to remove the largest fragments and concentrate the fine litter. Concentrated litter from each quadrat was then put in its own mini-Winkler sacks for 2 days in the lab. Winkler samplers are common and efficient for quantifying ant abundance and diversity (Fisher 2005). After 2 days, all worker ants were extracted and enumerated. The code to reproduce our analysis can be found at https://github.com/MoBiodiv/elev_gradient.

Overall, we found that species richness decreased with elevation at the spatial grain of the individual pitfall traps (Supplemental Fig. S1, β_1_ = −0.009, *R*^2^ = 0.59) but the decrease in total abundance at best was weak (Supplemental Fig. S1, β_1_ = −0.171, *R*^2^ = 0.10). Because of the strong effect of elevation on S, but weak effect on N, this suggests that it is unlikely that S is declining solely as a result of a sampling effect (i.e., the more-individuals hypothesis) along the elevational gradient. Note that OLS coefficients are fairly robust to the patterns of heteroscedasticity apparent in Supplemental Fig. S1.

To more fully decompose the components of diversity changing along the elevational gradient, we deployed the full multiscale MoB analysis using mobr (Fig. 4). The sSBRs show a general trend of higher S at lower elevations (Fig. 4A, darker curves), but the shape of these curves vary with spatial scale (x-axis). Note that many of the sSBRs cross at intermediate scales indicating that the ranking of site diversity across elevations depends on scale. The nsSBR, which ignore spatial aggregation, also tend to show that the lower elevation sites have higher S (Fig. 4B). Although again, many of these nsSBR curves cross at intermediate scales (Fig. 4B) indicating a signature of scale-dependence. The IBRs are qualitatively similar to the nsSBRs.

**Figure 4.**
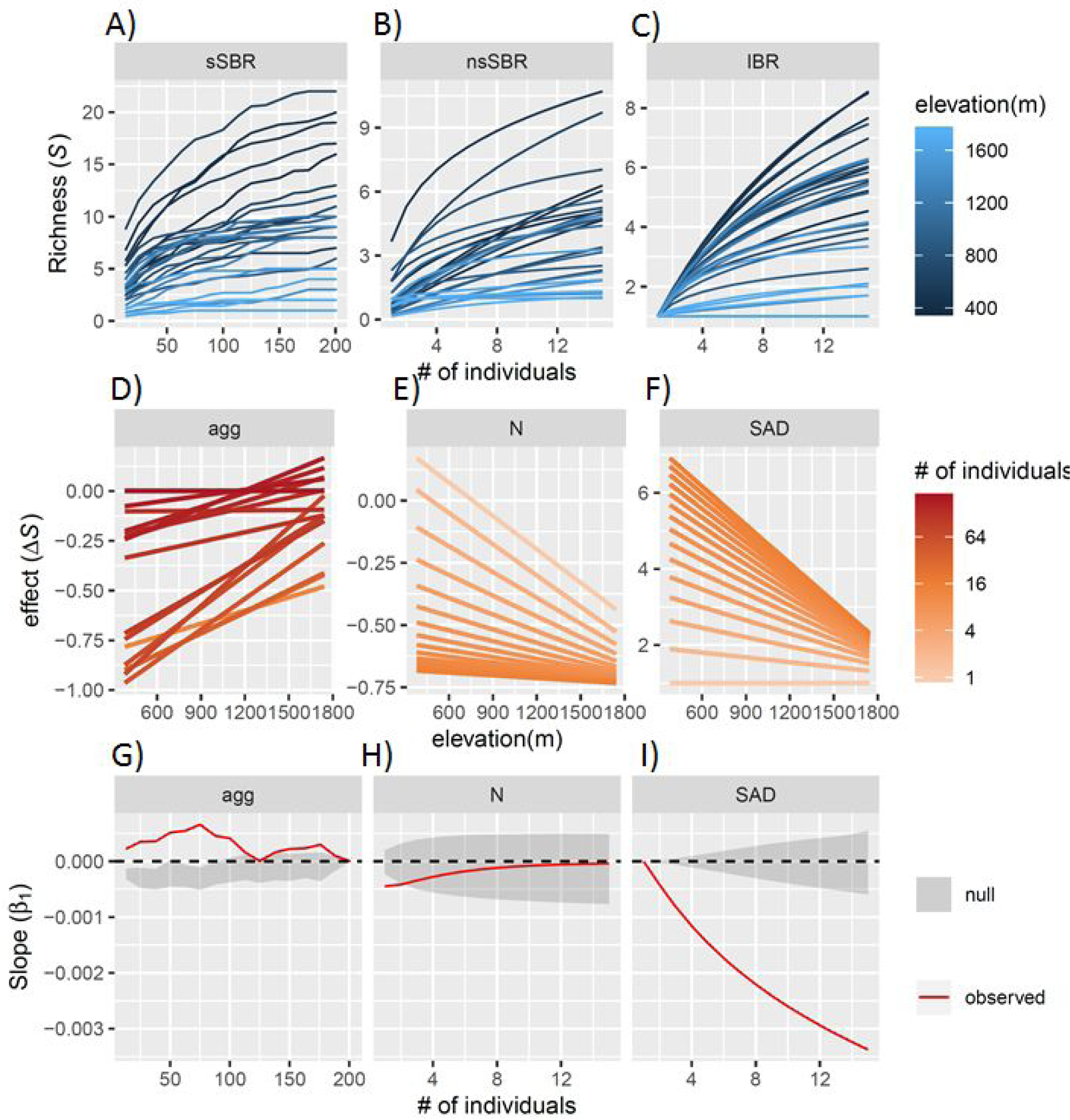
Multiscale analysis for the ant communities. A) The spatial, sample-based rarefaction (sSBR), B) the non-spatial, sample-based rarefaction (nsSBR), and C) the individual-based rarefaction (IBR) all expressed against number of individuals where each curve was constructed from a different site along the elevational gradient (black to blue lines). Panels D-F, show the regression lines of the linear model of ΔS ∼ elevation (m) at each sampling scale (light orange to dark orange lines) due to D) aggregation (agg), E) density (N), and F) evenness (SAD) effects. Note that the sampling effort color gradient is log transformed. Panels G-I, show how the OLS slope for each component of community structure changes across sampling efforts relative to null model expectations (grey polygon is the 95% quantile of the null model)

The change in species richness due to the aggregation effect tended to increase with elevation (Fig. 4D positive slopes), but this relationship weakened at larger sampling efforts (Fig. 4D, darker red lines have shallower positive slope). The N effect decreased along the gradient (negative sloping regression line) but this was only apparent at the finest spatial scale (light orange lines). The SAD effect also decreased with elevation but it became stronger with sampling effort (note how the regressions shift from flat light orange lines to negatively sloping dark orange lines).

The relationship between elevation and aggregation effect was more positive than expected under a null model of randomly distributed individuals (Fig. 4G), indicating species are less spatially aggregated at higher elevations. However, this effect was fairly weak: the magnitude of the largest slope was 0.0007 species per meter, which equates to about half a species across the approximately 1000 meters of elevation covered by the gradient and decreased as a function of sampling effort. The N effect did not differ from a random expectation at any scale except the very finest (Fig. 4H); therefore, we can conclude that low elevation sites are not more species rich because they have more individuals (which is consistent with Fig. 4). The slope of the SAD effect was negative along the elevational gradient, reaching a maximum of −0.0034 species per meter, (Fig. 4I). This indicates that higher elevation sites have lower S because of greater species dominance. Across the approximately 1000 m range in elevation we expect this effect to cause on average 3 fewer species to occur in a pitfall trap at a high elevation sites.

Using the multiscale MoB analysis we found that the Smokies ant diversity gradient is largely driven by changes in the SAD and aggregation effects. Interestingly, these two components of change have counteracting effects on *S* in this system. However, because the negative SAD effect is stronger than the positive aggregation effect, the net effect is that *S* decreases with elevation.

## Discussion

Diversity gradients are rich testing grounds for ecological theory. However, the most common metric of diversity, species richness, may respond similarly to different processes and thus cannot provide unambiguous tests. Our extension of the MoB analysis to continuous explanatory variables allows us to decompose diversity gradients into the effects of the different components--evenness, density or spatial aggregation--changing along the gradient. By describing the influence of changes in these components on changes in richness, we can provide more powerful tests of ecological hypotheses.

An example is the dataset on ants that we described above. One major feature that varies along elevational gradients is the amount of energy available to species. In species-energy theory, the more-individuals hypothesis (Wright 1983, Srivastava and Lawton 1998, Storch et al. 2018) proposes that richness should be driven by N effects. In the ant dataset we examined, however, we found no support for this hypothesis. Although there was a slight decrease in total ant abundance with increasing elevation (Fig. 4), this reduction in N did not drive the observed species richness pattern (Fig. 4). Instead, we found that richness declined primarily because higher elevation sites had lower evenness in their SAD, and to a lesser degree, because there was lower within-site aggregation of ant species at higher elevations. Many hypotheses can be linked to shifts in the SAD and spatial aggregation we observed (e.g., changes in competitive dominance, dispersal limitation, and/or environmental filtering) and information beyond what our analysis considers would be necessary to more fully differentiate these hypotheses.

More generally, decomposing richness into its components along ecological gradients may help provide resolution to apparently discordant empirical patterns of richness. For example, little consistency has emerged from some of the most well-studied ecological gradients of species richness, such as those along disturbance gradients (Mackey and Currie 2001, Svensson et al. 2012) and productivity gradients (e.g., Mittelbach et al. 2001, Adler et al. 2011). Some of the variation observed along these gradients is most certainly due to differences in the scales in which observations are taken (e.g., Rahbek 2005, Chase et al. 2018), but much of the variation could be due to the differential influence of these gradients on the components of species richness, such as on the density of individuals, the SAD or within-species aggregation. By examining how these components change along gradients in a more consistent way, we can begin to achieve greater synthesis than is currently possible with information only on species richness.

While the gradient version of MoB provides an important advance over the previous version that was only able to compare among categorical variables, there are many more directions in which the framework could be extended further. For example, MoB relies on data of individuals of each species so that rarefactions can be performed. Such data are often unavailable and only presence-absence data are available. For such cases, it should be straightforward to apply MoB to presence-absence data with a goal to partition changes in richness due to occupancy and spatial aggregation (see e.g., Tjorve *et al.*, 2008 for a similar approach using presence-absence data). However, for some taxa, separation into individuals is difficult if not impossible, and relative abundance data are instead available as estimates of visual cover or biomass. It is less clear how to interpret MoB metrics when using cover or biomass which in many communities may not be correlated with numbers of individuals. Finally, although we applied our approach using linear models of diversity change along a single explanatory variable (e.g., elevation). Logical next steps would be to consider a multiple regression framework in which the partial effects of several variables are considered simultaneously, as well as to include the potential for non-linear effects.

## Acknowledgments

A grant from Discover Life in America grants to N.J.S. supported collection of the empirical ant data. Jaime Ratchford, Raynelle Rino Melissa Geraghty, Chuck Parker and Keith Langdon facilitated this work in a variety of ways. The contributions of J.M.C, T.M.K., T.E, and S.A.B. were enabled by the German Centre for Integrative Biodiversity Research (iDiv) Halle-Jena-Leipzig funded by the German Research Foundation (FZT 118); the methods presented here emerged from several workshops funded with the support of iDiv (to JMC) as well as from the Alexander von Humboldt Foundation as part of the Alexander von Humboldt Professorship of TMK.

