## Supplemental Figure 1 for "A multiscale framework for disentangling the roles of evenness, density and aggregation on diversity gradients"

**Supplemental Figures**

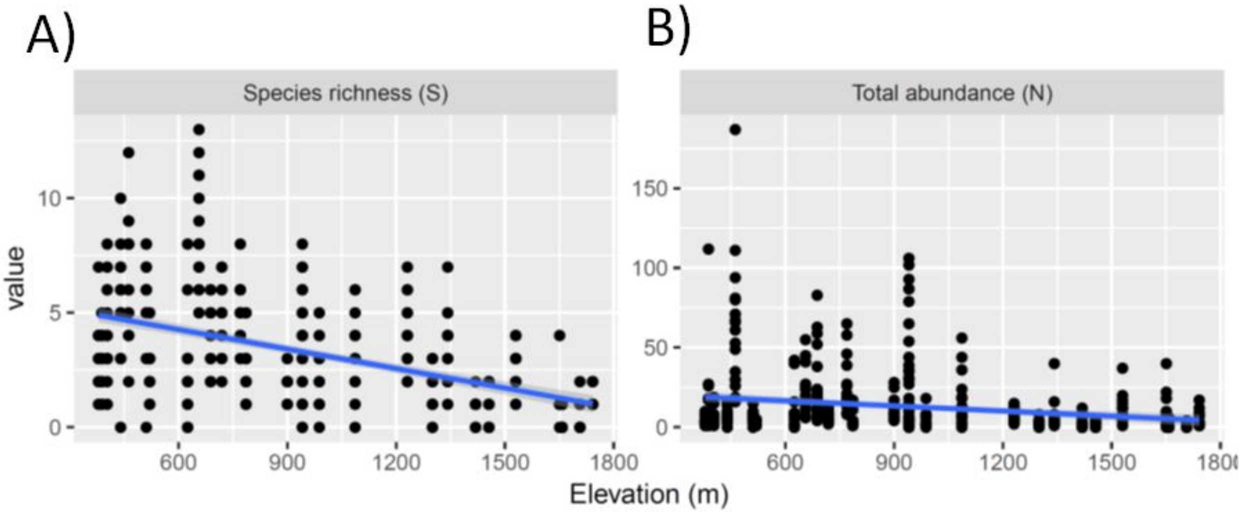


Fig. S1. Elevational gradient in (A) species richness (SW, $\beta_1$ = -0.009, $R^2$ = 0.59) and (B) total abundance (N, $\beta_1$ = -0.171, $R^2$ = 0.10) of the ant Winkler samples.
